# Polli-markers: spectral and chemical biomarkers for detecting cryptic early plant pollination responses

**DOI:** 10.64898/2025.12.03.692026

**Authors:** Catherine Parry, Colin Turnbull, Laura M.C. Barter, Michael Smith, Panagiotis Barmpoutis, Kamil Skirlo, Hannah V. Florance, Richard J. Gill

## Abstract

- Pollination is essential for plant reproduction, ecosystem resilience and human health. Yet, our capability to map pollination service delivery in real-time across large areas remains poor. Determining where and when flowers are pollinated is vital to mitigate widespread pollination deficits, increase plant health and yield, and support pollinator management. Hence, innovative approaches are urgently needed for establishing scalable predictive bioindicators of plant pollination status with the goal of achieving real-time landscape-scale monitoring.
- Here we present two parallel controlled pollination assays in which we characterise the post-pollination petal physiology of a world leading flowering crop, *Brassica napus*, using *in-situ* close-range hyperspectral reflectance and semi-untargeted metabolomics.
- This multiomics approach coupled with supervised machine learning and biomarker detection reveals cryptic changes in the UV petal reflectance spectrum which are predictive of pollination status, representing a novel set of candidate pollination bioindicators (‘*polli-markers’*), and our high-resolution time series enables prediction of when this pollination event occurred. It also reveals an associated set of candidate metabolites, including flavonoids and senescence markers, shedding light on the functional pathways related to our polli-markers.
- This study provides key insights into floral development, enabling a transformative step towards predicting, mapping and quantifying pollination service delivery at the landscape scale.

## Introduction

Vital for plant reproduction and health, pollination is a process underpinning trophic networks and the stability of ecosystems worldwide (Ollerton, 2020). For agricultural crops, pollination drives high yields making it an essential service for food security, human health and the global economy (Eilers *et al*., 2011; Breeze *et al*., 2016; Khalifa *et al*., 2021). But as animal pollinator populations decline and the human population grows, pollination demand is increasingly outstripping supply, making efforts to mitigate deficits an urgent priority (Webber *et al*., 2020; Olhnuud *et al*., 2022; Smith *et al*., 2022). To address this challenge, effective tools to accurately determine where and when pollination is delivered in real-time are needed (Parry *et al*., 2024). Only when armed with such information can we deploy targeted strategies to remedy pollination deficits, such as through pollinator supplementation and landscape management (Gill *et al*., 2016). Preventing us from reaching this goal is the lack of modern toolkits to reveal reliable biomarkers of plant pollination status at useful spatiotemporal scales (Breeze *et al*., 2021).

To date, estimating pollination delivery during a season has typically relied on anecdotally timed crop walks or observations of animal pollinator activity. These efforts require significant time investment, are expensive, and difficult to conduct at scale (Perrot *et al*., 2019; Hutchinson *et al*., 2022). Moreover, we cannot assume that measures of pollinator foraging directly translate to pollination success, especially in mass flowering species (Garibaldi *et al*., 2020; Page *et al*., 2021). If we are to improve mapping of pollination delivery, we need to identify biomarkers based on honest pollination signals to provide direct measures of pollination events. Coupled with remote sensing and/or sampling technologies, automated detection of pollination delivery in near-real-time could be achieved at high resolution and across suitable spatiotemporal scales (Parry *et al*., 2024). For instance, spectral reflectance measures have the potential to provide reliable assessments of plant syndromes like nutritional deficits and disease (Zhang *et al*., 2019; Gonzales *et al*., 2022). However, to our knowledge, such an approach is yet to be directly applied to pollination syndromes.

Flower petal-based biomarkers have the potential to enable the detection of rapid pollination-triggered signals whilst generating a canopy-scale overview of pollination, especially for mass flowering crops in agricultural settings. Furthermore, given it is the organ which directly interacts with a pollination vector (e.g. insects), we could eavesdrop on honest signals from an individual flower or plant to viable pollinators. We know petals to be transient structures, whose senescence may be premature or delayed in response to pollination (Rogers, 2006) with their fate determined by the regulation of senescence associated genes and, importantly, shifts in primary and secondary metabolites (Stead & Reid, 1990; Farzad *et al*., 2002; Tripathi & Tuteja, 2007). Senescent petals undergo significant changes in their biochemistry, giving rise to marked changes in cellular ultrastructure (Smith *et al*., 1992; Karabourniotis *et al*., 2021), properties which together have the potential to alter light refraction and absorption, measurable in its spectral reflectance profile (Steidle Neto *et al*., 2009). Such spectral changes, however, will be subtle and typically imperceptible to the human eye (Ohashi et al., 2015). This means that successful pollination events go undetected for days to weeks until petal drop and a seed pod begins to develop, making crop-pollinator management and conservation decisions challenging. Taking a multiomics approach, the aim of this study is to identify dual spectral and biochemical markers of plant pollination to reveal previously cryptic signatures of a pollination event directly after the event has occurred.

In taking the first steps towards field-functional ‘polli-markers’, our objectives were threefold: identify which regions of the spectral reflectance profile can act as polli-markers; ii) identify which candidate compounds in petals can act as pollination indicators; iii) assess if floral metabolites shed light on the functional pathways underlying identified spectral polli-markers. We conducted experimental assays on the globally leading, pollination-dependent flowering crop *Brassica napus* (Canola, oilseed rape) in which we performed tightly controlled pollination treatments to track changes in petals at high resolution (Fig. 1). First, we measured single-point hyperspectral reflectance and applied supervised machine learning (support vector machine (SVM)) to our spectral profiles to predict pollination status (pollinated vs. unpollinated) and recursive feature elimination (RFE) to detect spectral polli-markers. We further applied SVM-RFE to identify biomarkers of floral age in pollinated flowers as a predictor of when a pollination event occurred. We determined whether spectral shifts occur in the human visible or non-visible range, and whether these changes occur discretely or are represented by combinations of spectral bands. In a second mirrored assay, we applied semi-non targeted metabolomics by liquid chromatography quadrupole time-of-flight (LC/QTOF) mass spectrometry. We characterise time-series changes in the petal metabolome in response to pollination, both to identify groups of compounds which could act as chemical indicators and to help mechanistically explain the functional underpinnings of our spectral polli-markers.

**Figure 1.**
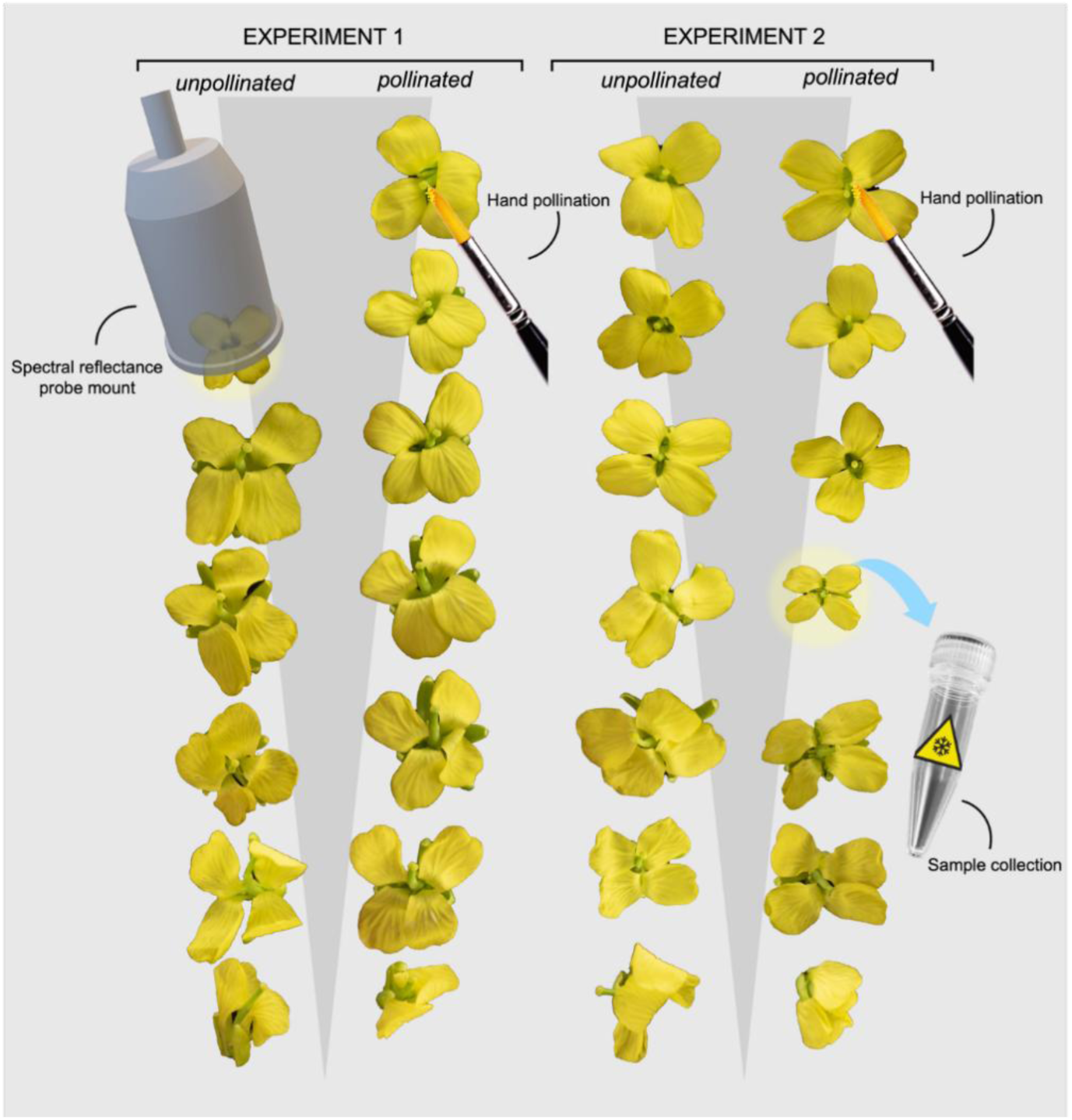
Measures taken in lab-controlled pollinated and unpollinated *B. napus* flowers in two time series experiments. All flowers were emasculated (anthers removed prior to anthesis) to prevent pollen contamination. *Experiment 1* depicts spectral reflectance measures taken *in situ* using the probe mount for illumination control, from flower opening in unpollinated flowers and, in the pollinated population, post hand-pollination. Repeat spectral reflectance measures were taken of each flower over the entire time-series, until petal drop. *Experiment 2* depicts destructive sample collection from pseudo-independent replicates from flower opening in unpollinated flowers and, in the pollinated population, post hand-pollination, whereby the flowers were immediately flash-frozen for metabolomic analysis. Sample collections were conducted to cover the complete time series. Images are not to scale.

## METHODS

### a. Plant material

*Brassica napus* cv. Westar (wild-type) seeds were grown in Levington F2+S medium and cold stratified for 4 days before being transferred to a separate controlled environment chamber under long day conditions (16-hour day under 120 μmol/m^2^ light intensity at 22°C, 8-hour night at 18°C with 60% humidity throughout). Once germinated, the seedlings were transplanted at the two true leaf stage into 9 cm diameter pots containing Levington M3 medium. Seedlings were then transplanted again when reaching the six true leaf stage into 11 litre extra-deep pots containing Levington M3. Plants were watered weekly, with two litres of Miracle-Gro fertiliser added to watering every two weeks until flowering.

### b. Experiment 1 – reflectance spectrometry experimental protocol

The experiment commenced as the first flower opened, and each flower to open was added to testing (range = 1-15 flowers per parent plant). Flowers were emasculated soon after opening, prior to anthesis, by removing the anthers with forceps. Additional plants acted as pollen donors and were allowed to flower freely in a separate controlled environment chamber to prevent pollen contamination. Upon opening, each flower was labelled with an ID ring to allow repeated measures of the same flower over its lifespan. Once the stigma became receptive, individual test flowers were pollinated by depositing pollen from the anthers of a pollen donor plant onto its stigma. Measures were taken every four hours during daylight hours, from the time of petal opening until the petals dropped (c.2 days). At the end of testing, flowers were allowed to develop freely until senescence, after which the seed pods from test and control plants were checked. Any seed pods developing on a control plant occurred due to pollen contamination and therefore the relevant IDs were excluded from data analysis during data cleaning.

### c. Experiment 2 – metabolomics experimental protocol

Whilst a separate experiment, the exact protocols used for assay 1 regarding experimental commencement, emasculation, pollen donation, assignment, flower ID, and controlled pollination method were again followed. But this time, after allotted time-periods, individual flowers were removed from the plant, placed whole in a cryovial and immediately flash-frozen in liquid nitrogen. The flowers were stored at -80°C until chemical extraction.

### d. Hyperspectral reflectance equipment

Hyperspectral reflectance measurements were taken from *B. napus* flower petals *in situ* using a QR600-7-SR125BX reflection/backscatter probe (Ocean Optics ©) connected to the Ocean Flame-SXR1 mini spectrometer, and data gathered using the Ocean View software. The petal tissue was illuminated with full spectrum light from a DH-2000_BAL Light Source (halogen and deuterium bulbs) (Ocean Optics ©). Spectrometer integration time was set to 1.59500E0 throughout, with electric dark correction enabled. External light (light other than that provided by the full-spectrum light source) was excluded using a bell-shape probe mount designed in Tinker-Cad and 3-D printed with matt black PLA filament (Supplementary Fig. 1). The probe was secured to the probe mount, and the mount placed over the flower *in-situ* with a high-absorbance black felt backing. The equipment was calibrated at the start of each testing occasion using a white standard (Ocean Optics ©). For each flower measurement, ten spectra were recorded in sequence and averaged (mean) during data processing.

### e. Spectral data preparation and pre-processing

All processing and analysis of hyperspectral reflectance data was carried out in R Studio (vers. 2023.12.0+369). In addition to base R, for data processing the following additional packages were used: ‘caret’ (Kuhn, 2008), ‘caTools’ (Tuszynski, 2024), ‘ChemoSpec’ (Hanson, 2024), ‘dplyr’ (Wickham *et al*., 2023), e1071 (Meyer *et al*., 2024b), kernlab (Karatzoglou *et al*., 2024), ‘stringr’ (Wickham, 2023), and ‘tidyr’ (Wickham et al., 2024)). For producing plots, the following additional packages were used: ‘ggnewscale’ (Campitelli, 2024), ‘ggplot2’ (Wickham, 2016), ‘ggrepel’ (Slowikowski, 2024), ‘ggspectra’ (Aphalop, 2015), and ‘patchwork’ (Pedersen, 2024). Data cleaning comprised trimming the spectra by 100 nm at each end to remove regions of high noise. Spectra were plotted to remove any erroneous measurements, and the first three principal components from a Principal Component Analysis (PCA) were visualised to identify further outliers, which were also removed (n = 2 erroneous spectra). To determine the most appropriate for maximising model performance, in addition to the raw data (cleaned but unnormalized spectra), data were normalised in each of three ways: a) probabilistic quotient normalisation, b) 1st derivative calculated with a Savitzky-Golay algorithm, and c) 2nd derivative calculated with a Savitzky-Golay algorithm. Data normalised with each method were used to train the SVM and the performance of each combination compared to determine the final model.

For model training the data were split into a training (75% of the total) and testing (25%) dataset in a stratified random manner, using a duplex algorithm to group by flower ID. For training each model, the training dataset itself was split in a stratified manner to account for repeat measures of individual flowers over time. During cross-validation we grouped by flower ID, with 75% (576 spectra) used for training and 25% (195 spectra) for validation in each iteration.

### f. Final model fitting and spectral biomarker detection

We applied support vector machine (SVM) models to predict pollination status (binary classification) and floral age in pollinated flowers (regression). We chose linear-kernel SVM to perform the analysis due to the dimensionality of the data, with our assumption that there will be little interaction between features, and that the two classes will likely be linearly separable. To explore whether the results depended on this choice, we applied a range of commonly utilised model types (see Supplementary Methods, Supplementary Fig. 8; 10). Parameters of the SVM (cost and gamma values) were tuned using a grid-search with 25-fold cross-validation. Data were scaled and mean-centred (Meyer *et al*., 2024a), and final models were executed with 10-fold cross-validation.

We conducted a permutation test to calibrate the significance of SVM models’ performance against a null distribution (Smit *et al*., 2007). The results of our permutation test for our binary classification (57%), and regression SVM (R^2^ = 0.02) reflect our slightly unbalanced dataset (Supplementary Table 1). We went on to test the impact of normalisation methods on performance for each respective SVM model.

Reducing dimensionality through feature selection not only reduces the computational load, but can also improve model performance, improve generalisability and increase the applicability of these findings (Zhang *et al*., 2006). RFE was therefore applied with the final models to determine the most important features (spectral wavelengths) for classification and define our biomarkers. We applied RFE with 6-fold cross-validation, retaining up to 170 features (10% of the total 1700 wavelengths) to simplify the model and maximise computational efficiency. The most informative features were extracted and considered as candidate biomarkers. Indeed, machine learning combined with RFE has been a successful approach to biomarker detection in many omics applications, as well as to hyperspectral reflectance datasets (Zhang *et al*., 2006; Yoosefzadeh-Najafabadi *et al*., 2021).

### g. Chemical extraction protocol

Frozen flower samples were freeze-dried for 24 hours, then petals dissected from the pistil by gently removing with forceps. The petal tissue was macerated for 90 seconds at 30 Hz in a tissue lyser (pre-chilled with dry ice) with a 1 mm steel grinding ball. Extraction buffer solution consisting of methanol/water/acetic acid 85:15:0.05 (v/v/v) was added at 1 ml per 100 mg petal sample then incubated at 4°C overnight. Samples were centrifuged at 10,000 g for 10 minutes and stored at -20°C until the day of analysis. Each sample’s supernatant was filtered separately through a 0.22 µm nitrocellulose syringe filter then 50 µl of each sample was aliquoted into a single well of a microplate (Thermo Fisher Scientific), with position of the samples in the microplate configured according to a randomised plate plan. A quality control (QC) sample was prepared by mixing 5 µl of extract from each individual and placed in every 10th well of the microplate.

### h. LC Q-TOF for metabolomics

For detection of metabolites, Quadropole Time of Flight LC/MS was applied. Agilent 1290 Infinity II UHPLC was coupled with Agilent 6546 LC/QTOF and equipped with an Agilent Poroshell C18 column (120 EC-C18 4/6×100 mm, 2.7 µm particle size). The mobile phases consisted of Solvent A (0.1% formic acid in water) and Solvent B (0.1% formic acid in Methanol). For each injection, 1 µl of sample was loaded, and the following gradient profile was applied: 5% B at 1 min, 100% B at 10.0 min, 100% B at 12.0 min, 5% B at 12.1 min, and 5% B at 15.0 min. The source parameters were set as follows: Drying Gas temperature at 320°C, Drying Gas Flow at 12 L/min, Nebuliser pressure at 40 psi, Sheath Gas temperature at 350°C, Sheath Gas Flow at 12 L/min, Nozzle Voltage at 3500 V, Fragmentor Voltage at 120 V, Skimmer Voltage at 65 V and Octuple Voltage at 750 V. Data was acquired in the m/z 50-1700 range in both positive and negative ion mode. The data was acquired using Mass Hunter Acquisition (version 11).

### i. Chemical library preparation and feature selection

A library was constructed from a literature search of metabolomics studies on flowers and petal development of Brassicaceae species (see Supplementary Methods for the list of publications used to compile the library). Mass and retention times were derived from Metlin (Guijas *et al*., 2018). The library was refined to 600 unique compounds based on a search of compounds present in a subset of six QC samples (evenly spread over the run time, including both positive and negative ion modes) using the Find by Formula algorithm in the Agilent MassHunter Qualitative software (version 10). Time alignment was then applied to all the spectra according to the middlemost QC sample. Feature selection was conducted in the Agilent Profinder software, and compound identities were confirmed manually based on a detection frequency (minimum presence in 20 samples), confidence score (threshold: 75%), and visual inspection of m/z tolerance and consistency of isotope distribution pattern.

### j. Cleaning, pre-processing and analysis of metabolomic data

Analysis was conducted with Metaboanalyst (Pang *et al*., 2024), using the online platform (version 6) and R Studio (Metaboanalyst version 4). A preliminary analysis of the data highlighted three highly abundant compounds to be skewing the distribution of the data by inducing a bimodality which showed no relation to treatment or any technical variables. These compounds (*is 3-sophoroside-7-glucoside, fumaric acid and coniferin*) were removed from the subsequent analysis. Missing values were imputed with the median value of each feature column, and any feature with more than 75% missing values were removed. The data were filtered according to their reliability (exceeding 25% relative standard deviation), variance (greater than 20% standard deviation), and abundance (10% of median intensity value). The data were normalised to the sample median, transformed to a base 2 logarithm, and Pareto scaling applied (van den Berg *et al*., 2006). Metabolite classes were assigned using ClassyFireR for hierarchical chemical classification based on ChemOnt taxonomy (Djoumbou Feunang *et al*., 2016).

To account for the effects of time and pollination treatment, analysis addressed two questions: i) is the petal metabolome associated with pollination treatment, using partial least-squares discriminant analysis (PLS-DA; with 10-times leave-one-out cross-validation) considering pollinated and unpollinated flowers (binary); ii) is the petal metabolome associated with floral development (age/time since the flower opened), using Pearson’s correlation with time since flower opening (0-32 hours) as a continuous variable, and pollination status as a categorical variable. Based on variables of importance in prediction (VIP) score, and Pearson’s correlation coefficient, respectively, as well as informed by taxonomical classification and putative biological function, we tested individual compounds’ associations with time and treatment we fitted linear mixed effects regression (LMER), with the interaction term of time * treatment and plant ID as a random effect to account for pseudo-replication. To test individual compounds’ associations with time only, we applied LMER with time as a fixed effect and plant ID as a random effect.

## Results & Discussion

### a. Spectral polli-markers: defining the machine learning approach and biomarker detection

Repeat-measures of spectral reflectance profiles were recorded from 80 hand-pollinated and 63 unpollinated flowers (from 15 parent plants; Supplementary Table 1), with pollination treatment validated prior to data analysis (see Methods). Each profile comprised of 1,700 frequency values ranging across the spectra from 216 to 925 nm, with a frequency resolution of 0.418 wavelengths/point.

We applied a binary SVM to interrogate the petal spectral reflectance profile as the first step in our biomarker detection workflow (Supplementary Fig. 1 for an illustrative summary). To optimise model performance, we defined the parameters by tuning the cost using a grid search (cost = 0.25), with data scaled and mean-centred (Meyer *et al*., 2024a). We tested the impact of different pre-processing methods on SVM performance, determined primarily by area under the curve (AUC) of the receiver operating characteristic (ROC), and model test accuracy (Figure 2A), which revealed raw data was highest performing (AUC = 0.7, accuracy = 70.25%). Multiplicative scatter correction (MSC) negatively impacted performance compared to the raw data, while first and second derivatives of the spectra had relatively small impacts on model performance. The regions of highest separability between the classes may therefore be co-located with low-frequency components rather than at a peak. The SVM was thus trained on raw data before being challenged against the test dataset with stratified cross-validation, achieving a test accuracy of 71.28% (Figure 4B). The model tended to correctly classify pollinated flowers more frequently than unpollinated, and performance substantially outperformed the classification accuracy of the permutation test (see Methods; Supplementary Fig.2).

**Figure 2.**
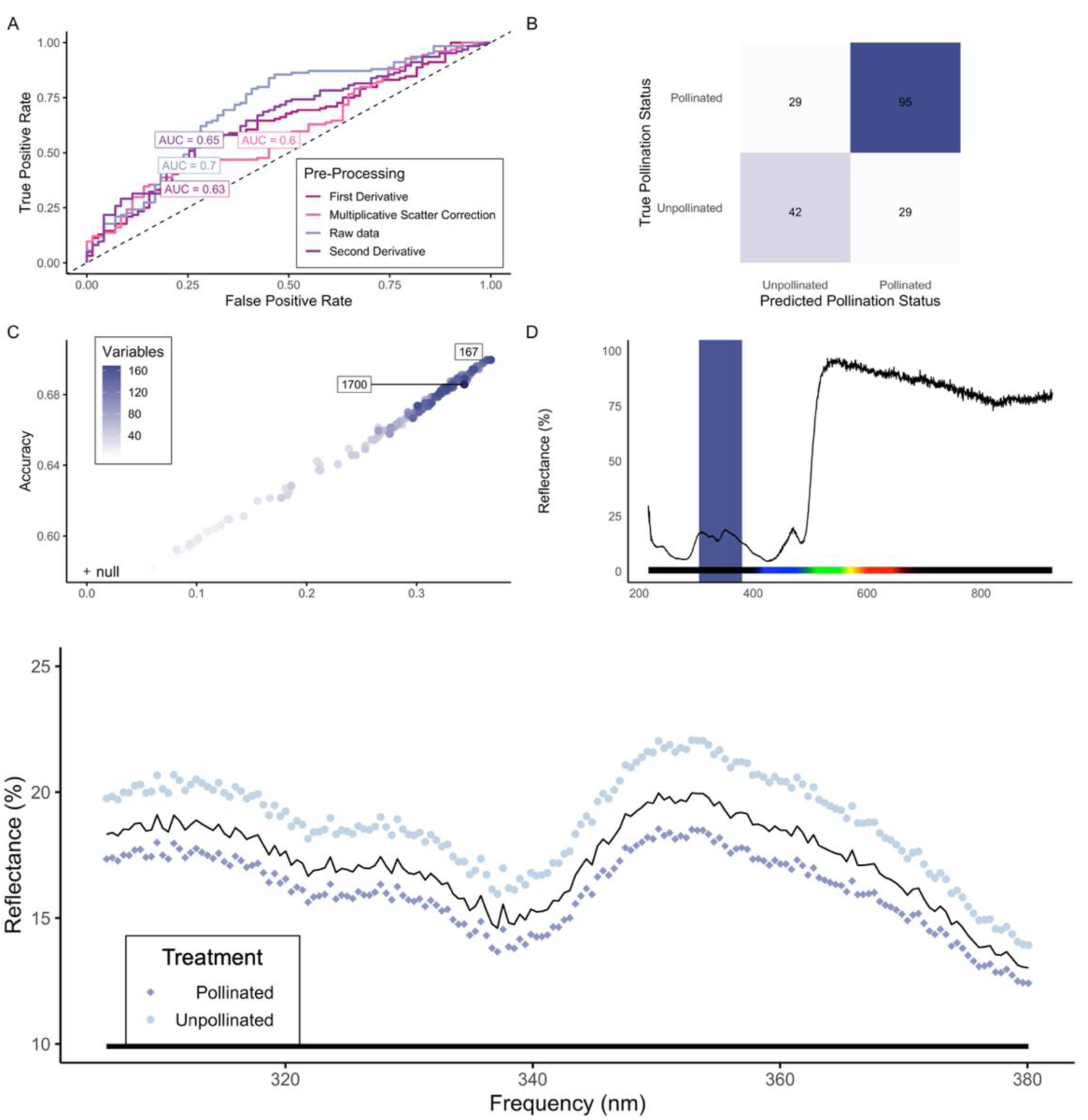
Identifying spectral polli-markers. A) ROC curves for SVM models trained and tested on pre-processed data. B) Confusion matrix of SVM model performance trained and tested on raw data to predict floral pollination status from spectral reflectance profiles. C) Performance of the SVM model under recursive feature elimination (RFE) to determine the optimal number of spectral wavelengths; ‘null’ indicates the model’s performance under a null distribution determined by a permutation test. D) Candidate spectral polli-markers determined by SVM and RFE highlighted against the mean reflectance spectrum. E) Spectra at the spectral polli-marker region; mean reflectance intensity in pollinated and unpollinated flower petals, with the mean spectrum across all flower samples regardless of treatment indicated by the solid black line.

We applied RFE for feature selection as a biomarker discovery tool. By ranking the coefficients and recursively removing the least informative features, RFE determined which of the total 1,700 spectral wavelengths (‘variables’) were the most informative for predicting pollination status (Figure 4C). Here, 167 variables were optimal, and had equivalent performance to the model utilising the full complement of wavelengths (accuracy = 69.96%). The variables considered as our putative spectral polli-markers in *B. napus* clustered in the UV region of the spectrum in a single band at 305.548 - 380.083 nm (Figure 4D) with a decrease in reflectance over the surface of the petal rapidly after the pollination event (Figure 4E). In addressing our hypotheses, this shows that our spectral polli-markers are represented as a change at a discrete region of the non-visible [to humans] part of the reflectance spectrum.

### b. Chemical polli-markers: metabolite analysis and identifying candidate chemical signatures

The experimental samples consisted of 26 parental plants. From these there were 70 unpollinated flowers (from seven parental plants), 146 pollinated flowers (from 19 parental plants), and 11 flowers sampled prior to opening. The cleaned metabolite data consisted of 514 compounds identified across the petal samples. The most abundant compound classes were organooxygen compounds, carboxylic acids and derivatives, and flavonoids (Supplementary Fig. 3).

Firstly, regardless of the pollination treatment, principal component analysis (PCA; Fig. 3B) indicated a continual shift in the metabolome across all samples, with a gradient of change as the flowers age. Indeed, our unopened flowers were most similar to the youngest flowers which were sampled 4 hours after opening. This observed trend with age aligns with our expectation that petals would senesce over the experimental assay, even if this senescence is largely undetectable with the naked eye. The gradient is reflected in the heatmap, where there appears to be a generally linear relationship between individual metabolite concentrations and time (Fig. 3A).

**Figure 3.**
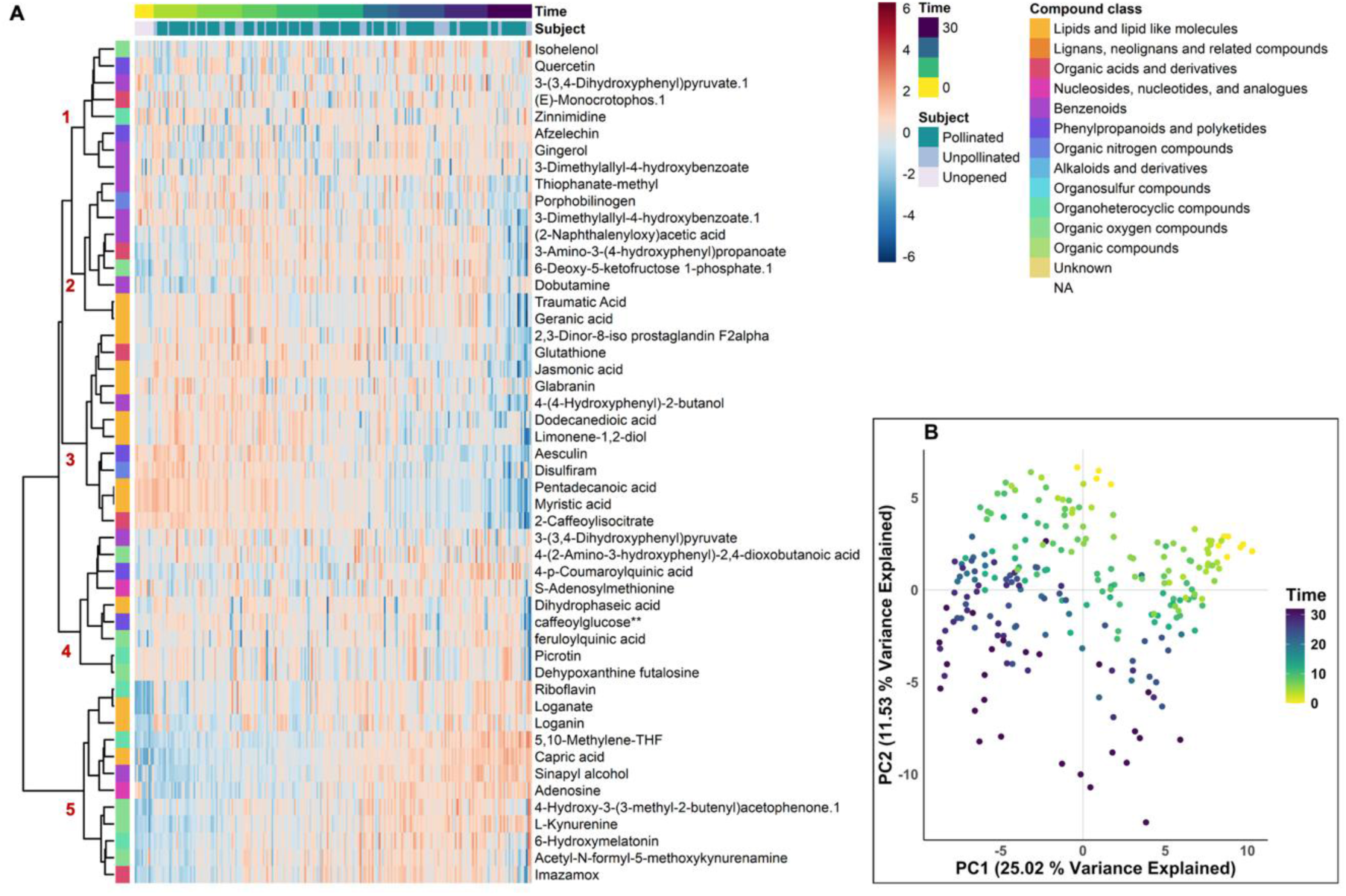
Metabolomic profiles of pollinated and unpollinated flowers in time-series. A) The heatmap of normalised compound concentrations (top 50 features) grouped by their pollination status and time. Compounds are clustered by Euclidean distance with a Ward algorithm and labelled by their taxonomic superclass. Groups 1-5 mark five major clusters which indicate correlations in compound occurrence. B) The first two principal components of PCA, with time indicated by colour intensity, 0 being unopened flowers.

**Figure 4.**
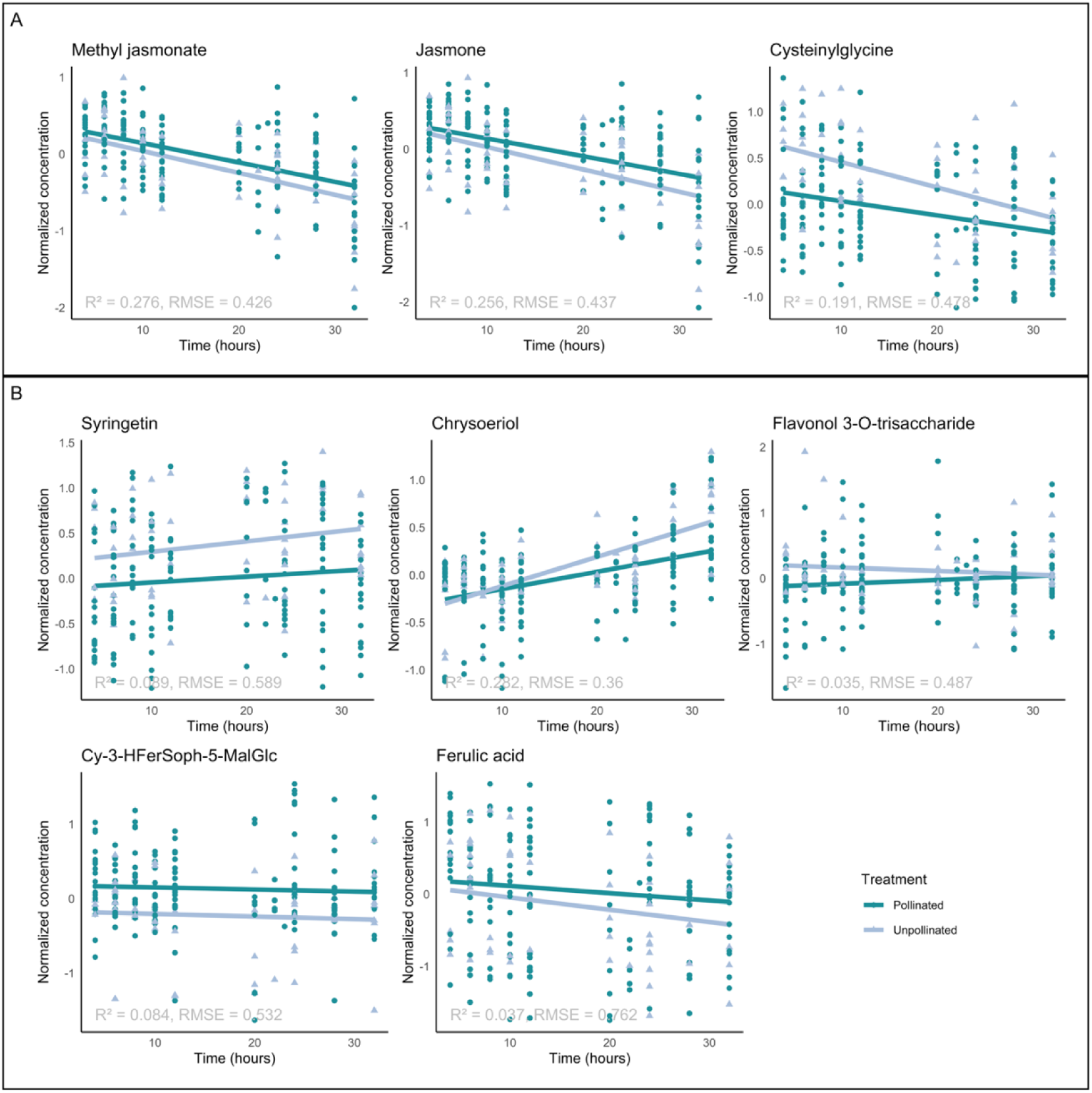
Temporal trends for normalised concentrations of metabolites that scored a high-VIP (biologically notable compounds) and thus represent candidates that independently or in combination can distinguish between pollination treatments. Metabolites are of A: putative growth-regulatory function, and B: phenylpropanoids and polyketides with putative pigmentation-related functions. Normalised concentrations plotted, with coloured lines which indicate the model estimates of treatment * time by a fitted linear mixed effects regression model (LMER).

Clustering of the metabolites indicates five major responsive groups, which tend to also be associated with their chemotaxonomical classifications (i.e. classes of chemicals that appear sensitive to time and treatment). Compound abundance of metabolites in the petals in group 2 and 3 (Fig. 3A), which are comprised of Benzenoids, phenylpropanoids, lipids and lignans, generally decline as the flower ages. Group 5 is dominant in organoheterocyclic and organic oxygen compounds, whose abundances in petals generally increase as flowers age. Concentrations of other individual metabolites, particularly group 1 and 4, appear to fluctuate over time.

Partial least squares discriminant analysis (pollinated vs. unpollinated flowers) highlighted 158 compounds to have variable importance in prediction (VIP) scores >1. The most important differentially occurring compounds (VIP >1.5) were dominated by compounds of benzenoid, lipid, organic acid, organic oxygen, and phenylpropanoid classes (Supplementary Fig. 4). The high-VIP compounds highlight two compound groups of biological interest: those with putative growth regulatory functions, and those categorised as flavonoids or flavonoid precursors (of class phenylpropanoids and polyketides; Fig. 4). We conducted a more conservative analysis of these high-VIP, biologically notable candidates (syringetin, Cy 3-hydroferuloylsophoroside-5-malonylglucoside, cysteinylglycine, Ferulic acid, jasmone, methyl jasmonate, chrysoeriol, flavonol 3-O-trisaccharide), looking independently at each individual compounds’ association with time as a function of pollination treatment (interactive effect). We fitted mixed effects models (LMER), with parent plant ID as a random effect to account for pseudo-replication. Whilst less sensitive to subtle patterns and not accounting for correlated or interacting groups of metabolites, this analysis did reveal differences between pollination treatment in this subset of compounds.

Cysteinylglycine declined in concentration over time (*p* = 1.22 × 10^−4^), with higher concentration in unpollinated flowers (*p* = 2.85×10^−4^, *q* = 2.23×10^−3^ (Benjamini-Hochberg False Discovery Rate); interaction effect *p* = 0.09, *q* = 0.24). Cysteinylglycine concentration profiles converge during the time series, suggesting an initial acute and rapid change in concentration within four hours of a pollination event. In addition to cysteinylglycine, other members of the glutathione metabolism pathway including ascorbate, L-cysteine and glutathione disulfide were also present in our samples, although were not associated with pollination treatment (see Supplementary Fig. 4). The flavonoid, Chrysoeriol, increased in concentration as flowers aged, and was significantly associated with pollination treatment and time (interaction *p* = 0.03, *q* = 0.23). Other phenylpropanoids and polyketides were consistent in concentration as flowers age, with little association with time, but several were different between pollination treatment. Flavonol 3-O-trisaccharide had relatively lower concentration in pollinated flowers (treatment effect *p* = 1.77×10^−2^, *q* = 0.07). Cy 3-hydroferuloylsophoroside-5-malonylglucoside is an anthocyanin whose concentration was elevated in pollinated petals (*p* = 2.8×10^−2^, *q* = 0.07), with a non-significant treatment effect, which may indicate an initial concentration shift occurred prior to the sampling window.

This indicates that whilst some metabolite changes do independently show strong association with pollination treatment, our analyses indicates that it is combinations of metabolites that better discriminates pollination status (by PLS-DA). Given the biological functions of these high-VIP metabolites, it is plausible that the combined effects of pathways associated with i) growth regulators, and ii) flavonoid composition contribute to discriminating pollination treatment, and our measured shift in the UV reflectance spectrum.

### c. Spectral biomarkers of floral age post-pollination

To estimate floral age post-pollination, we applied a SVM regression model (Radial Basis Function (RBF) kernel), and defined the parameters (cost = 2, gamma = 0.00075) with data scaled and mean-centred. We first tested the impact of pre-processing on SVM performance, determined by AUC of the regression error characteristic (REC), root mean squared error (RMSE), and R^2^ (Figure 5A). Model performance was similar across all pre-processing methods. The MSC-trained model’s REC AUC was higher (0.79) than the raw data (0.75), however performance of the model trained on raw data was overall superior (RMSE = 9.11 hours, R^2^ = 0.65). First and second derivatives of the spectra had little impact on model performance, again suggesting the regions of highest separability between the classes are co-located with low-frequency components rather than at a peak. Scatter-correction also had little impact, perhaps due to the relatively low degree of noise in our dataset measured under controlled conditions.

**Figure 5.**
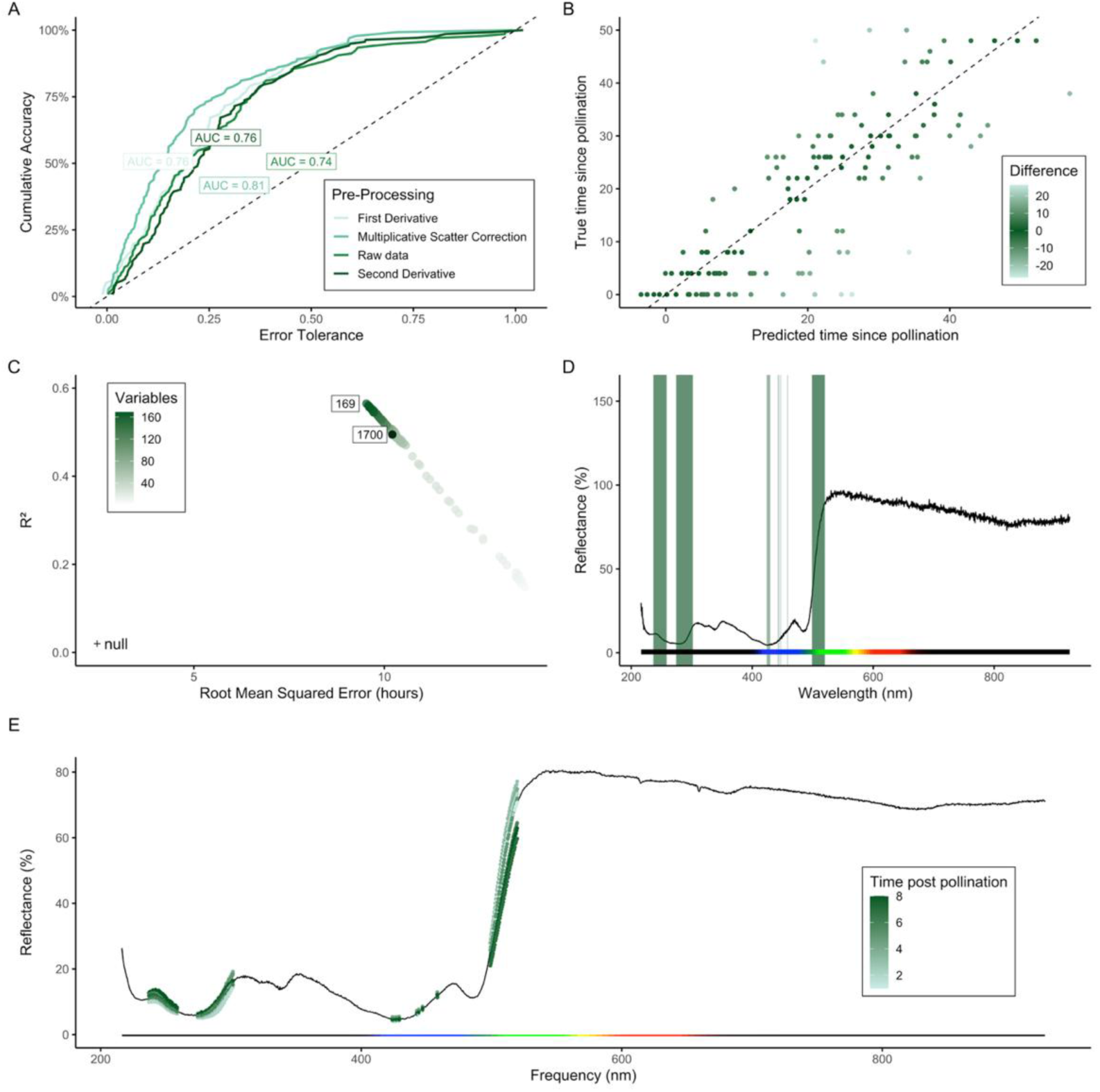
Determining spectral biomarkers of floral age post-pollination. A) REC curves for SVM models trained and tested on pre-processed data; the dashed line represents null/ accuracy under random prediction. B) Predicted vs. True (observed) time since pollination, with the dashed line this time representing perfect prediction. C) Performance of the SVM model under recursive feature elimination to determine the optimal number of spectral wavelengths; ‘null’ indicates the model’s performance under a null distribution determined by a permutation test. D) Candidate spectral biomarkers of floral age, determined by SVM and RFE highlighted against the mean reflectance spectrum. E) Spectra at the candidate biomarker region; mean reflectance intensity over time, with the mean spectrum across all flower samples indicated by the continuous black line.

The SVM trained on data normalised by MSC was challenged against the test dataset. This model achieved an R^2^ of 0.65 and RMSE of 9.11 hours; predicted vs. actual time since pollination is visualised in Figure 5B. The error rate in predicting flower age up to 12 hours after a flower received pollen was higher (RMSE = 8.67) but reduced in flowers 15-30 hours after receiving pollen (RMSE = 6.91). Pre-pollination measures (0 hours since pollination) appear to be outliers, with a high error rate of prediction by our model, but by approximately 15 hours post-pollination the floral age can be predicted with greater certainty.

The RFE resolved 169 wavelengths as optimal (Figure 5C) with this model predicting time since pollination with R^2^ of 0.56 (RMSE = 9.53 hours), higher than the model utilising the full complement of wavelengths (R^2^ = 0.49). These wavelengths are clustered primarily in the UV and green region of the spectrum, ranging between 237.009 - 256.77 nm, 274.6 - 301.467 nm, and 499.967 - 519.700 nm, and twelve discrete bands between 423.859 - 458.42 nm. We consider these 169 wavelengths to be biomarkers of floral age in pollinated flowers and thus could be applied as complementary predictors of when a pollination event occurred (Figure 5D). Petal average reflectance in the UV region (237-301 nm) shows evidence of an increase in intensity as the flower ages post-pollination, while concurrently reflectance subtly declines in the visible green region (499-519 nm; Figure 5E).

### c. Chemical biomarkers of floral age post pollination

Considering the change in metabolite concentration over time, 46 metabolites had a Pearson’s correlation coefficient greater than +/- 0.65 (Figure 6; Supplementary Fig. 6). These comprise metabolites from a diversity of classes: benzenoids; lignans, neolignans and related compounds; lipids and lipid like molecules; nucleosides, nucleotides, and analogues; organic acids and derivatives; organic compounds; organic oxygen compounds; organoheterocyclic compounds; and phenylpropanoids and polyketides. Ten of the most correlated metabolites are lipids or lipid-like, six of which showed a decline in concentration over time and four an increase, following a linear relationship (Figure 6). Of the metabolites most correlated with time, there are two notable groups; i) long chain saturated fatty acids, including myristoleic acid (C14:1), pentadecanoic acid (C15:0) and myristic acid (C14:0), which are the most strongly down-regulated metabolites (Fig. 6A), and ii) medium-chain saturated fatty acids, including Caprylic acid (C8:0) and capric acid (C10:0), which are the most strongly up-regulated metabolites (Fig 6B).

**Figure 6.**
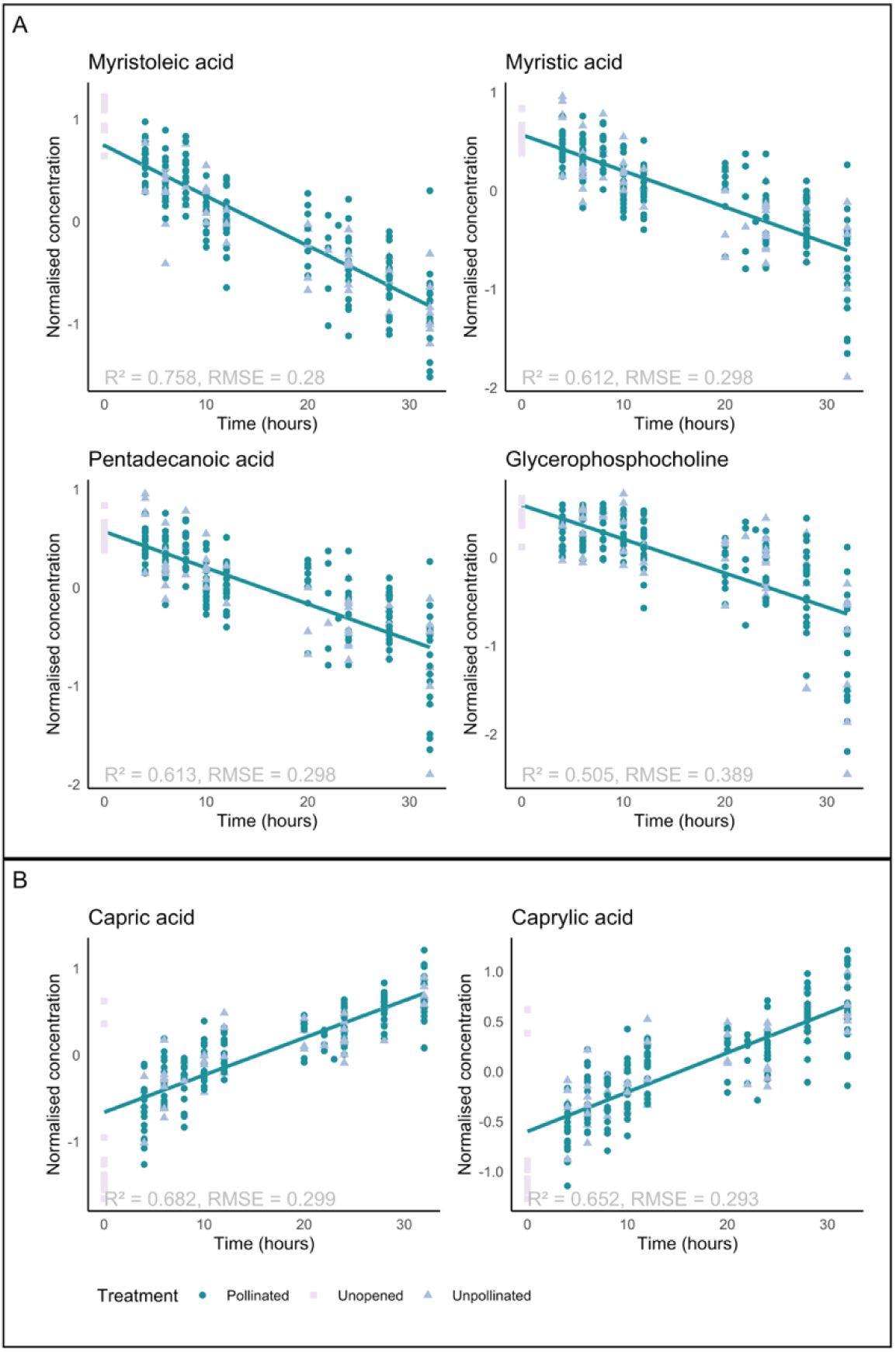
Temporal trends for normalised concentrations of focal metabolites. Focal metabolites scored a high-VIP (biologically notable compounds) as candidates for predicting floral age (i.e. concentration strongly corelates with time independently of pollination treatment). Compounds are of the lipid and lipid-like class, with group A fatty acids downregulated over time and B fatty acids upregulated over time. Coloured line indicates average prediction of floral age (time) by a linear mixed effects regression (LMER).

Our hyperspectral reflectance profiles reveal previously undetected cryptic changes in the *B. napus* phenotype in response to pollination. A rapid decrease in average UV-spectral reflectance intensity from the petal surface of pollinated flowers represents our candidate spectral polli-markers. In parallel, our semi-non targeted metabolomic study highlights a dual set of petal metabolites whose concentrations shift in response to pollination. In particular, variations in pigment-related metabolites, namely flavonoids, and metabolites related to growth regulation, are correlated with pollination status. Our chemical polli-markers are metabolites which putatively underly our spectral polli-markers, and this multi-omic approach presents candidates to be further explored and developed for a definitive and field applicable multimodal pollination bioindicator.

Crucially, our spectral polli-marker aligns closely with the spectral absorbance properties of the flavonoids (300-380 nm, Taniguchi et al., 2023), which we highlight as chemical polli-marker candidates. This suggests a compelling mechanistic link between the chemical and spectral indicators that we detect, suggesting (in this controlled setting,) our spectral polli-markers are representative of a cryptic phenotypic pollination response. Flavonoid metabolism has been shown to exhibit a high degree of plasticity in response to environmental stimuli and development, mediated by differential expression, transcriptional- and post-transcriptional regulation (Dong & Lin, 2021; Stavenga *et al*., 2021; Koski *et al*., 2024). Moreover, the structure of flavonoid molecules and their modifications strongly influence their reflectance spectra (Liu *et al*., 2021; Taniguchi *et al*., 2023) and are thus strong candidates for functional bioindicators.

In addition to pigmentation change, we highlight metabolites from across class groups with putative growth regulatory functions which are associated with pollinations status. Jasmonic acid is key for floral development, as well as important for pollination and stress responses (Yuan & Zhang, 2015). Jasmonate has been shown to be required for limiting cell expansion and thus ultimate petal organ size in *Arabidopsis* (Brioudes *et al*., 2009), and (exogenous) methyl jasmonate to promote flowering in *B. napus* (Pak *et al*., 2009). Jasmonic acid signalling also modulates floral volatile production and mediates floral attractiveness to pollinators (Stitz *et al*., 2014; Li *et al*., 2018). Conversely, cysteinylglycine is elevated in unpollinated flowers soon after opening, with concentrations declining as the flower ages. Cysteinylglycine is a degradation product of glutathione, an essential coenzyme and key reactive oxygen species scavenger which has important roles in stress and senescence responses (Cairns *et al*., 2006; Noctor *et al*., 2012). Elevated cysteinylglycine levels indicate active cell turnover, while comparatively depressed cysteinylglycine levels in pollinated flowers suggest ageing is accelerated from as early as 4 hours post-pollination (Ding *et al*., 2016). By 32-hours the concentrations of cysteinylglycine in pollinated and unpollinated flowers converge as senescence processes are also triggered in unpollinated flowers. Our results suggest the combined effect of jasmonates and other putative growth regulators (5-hydroxyindoleacetylglycine, picrotin, see Supplementary Fig. 6) with suppression of cysteinylglycine having a role in stimulating development in pollinated flowers, ultimately culminating in accelerated senescence.

Our pollination signals appear distinct from those associated with senescence or floral development. While the spectra of pollinated flowers do shift during development, these pollination-induced senescence signals are distributed across the UV and visible spectrum (Figure 5). Changes in spectral reflectance are subtle, but our biomarker detection pipeline reveals combinations of wavelengths which are important for predicting floral age post-pollination. The metabolomic markers most strongly associated with floral age are lipids or lipid-like, six of which decline and four increase in concentration over time in a linear fashion. Lipids have diverse functions, and given the dynamic nature of lipid organisation can act as useful biomarkers for environmental stimuli (Reszczyńska & Hanaka, 2020). The most strongly down-regulated metabolites are myristoleic acid, pentadecanoic acid and myristic acid, which are long-chain fatty acyls, and glycerophosphocholine which is associated with membrane turnover and phospholipid degradation (Van der Rest *et al*., 2002). Interestingly, two of the most up- regulated compounds are Caprylic acid and Capric acid, which are medium chain fatty acyls.

The dynamic up- and down-regulation of lipid components suggests a rebalancing of fatty acid composition over time, with an overall decline in long-chain fatty acids and phospholipid components, concurrent with a relative increase in medium-chain fatty acid components which may represent degradation products (Thompson *et al*., 1998). Dysregulation of fatty acid synthesis is a known marker of senescence, and degradation and reduced biosynthesis of fatty acids has been observed in other senescing tissues (Wagstaff *et al*., 2003; Yang & Ohlrogge, 2009). Interestingly, our most correlated metabolites change in a continual and directional manner throughout floral ageing, suggesting lipid remodelling processes commence soon after flower opening. This is in line with reports in *Arabidopsis* which show changes in cuticular fatty acid content to be linear during development from flower opening to reclosing (Alexander *et al*., 2021). The association between dynamic lipid restructuring and spectral shifts during development suggests cuticle features, and possibly cell shape, play a role in spectral scattering in the green region, resulting in the localisation of our spectral biomarkers region (Camarillo-Castillo *et al*., 2021).

A significant proportion of colour-changing flower species exhibit a change in spectral reflectance during their development, without a perceptible colour change (Ohashi *et al*., 2015). To our knowledge, however, this is the first demonstration of a strictly pollination-induced differential spectral signal, and indeed, the first demonstration of an imperceptible reflectance shift in an important agricultural crop. Our measured shift in UV reflectance overlaps with the visual sensitivity of many pollinators, in particular bees (Chittka & Menzel, 1992). Bees have a foraging preference for relatively higher UV reflectance (Klomberg *et al*., 2019) and are sensitive to small shifts in pigment concentration, therefore it is plausible that our spectral polli-markers reveal an honest signalling strategy (Papiorek *et al*., 2013). The UV shift we measure is subtle, yet it is uncertain how small changes in the spectral quality of a flower individual may be perceived over the canopy scale. This should be of particular consideration for mass flowering monocultures such as *B. napus*, wherein the canopy (a mixture of pollinated and unpollinated flowers across developmental stages) provides a long-distance signal, and individual flowers may signal pollination status at close range.

Increasing experimental throughput will no doubt be valuable to maximise model training accuracy, and expand our understanding of the context dependency of such molecular and biochemical pathways. We must apply multispectral indices based on our biomarkers in controlled and field settings, as well as determining whether our floral biomarkers are effective across cultivars, and indeed relevant across species. Exploring the application of our bioindicators across taxa and environmental conditions will be essential to answering these questions. As field-tested bioindicators are established, these must be integrated with remote sensing technologies to bring this novel pollination monitoring tool into the hands of farmers and landscape managers (Parry et al., 2024). Such a tool enabling field- and landscape-scale pollination profiling would facilitate the generation of unprecedented pollination datasets, filling knowledge gaps in pollination monitoring, plant-pollinator networks and ecosystem health. Our polli-markers are the first step to achieving a transformative and essential monitoring technology for mapping pollination across landscapes.

## Acknowledgements

We thank Paul Beasley, Po-Yuan Shih and Lisa Haigh for technical support, and Oliver Windram, Chris Adams, Austin Burt, Will Pearse, Manel Benlloch Guajardo and Matthew Tyler for inspiration and feedback on ideas. Brassica seed was provided by Graham Teakle and Lauren Chappell at the Warwick Crop Centre under MTA. Mass spectrometry data for metabolomics research was acquired at the Agilent Measurement Suite research facility at the Imperial College White City Campus.

## Contributions

RJG conceived the project, CP & RJG developed the concept. CP & RJG designed the spectral reflectance experiments with advice from CT and LB; CP & RJG designed the metabolomics experiments with advice from CT, LB, KS & HF. CP conducted the experiments, with support from KS for the metabolomics. CP conducted analyses with guidance from RJG, CT, MS, PB and HF. CP and RJG wrote the first versions of the manuscript, with final inputs from all authors.

## Supplementary Information

### Supplementary Methods

#### Testing protocol

Once the stigma became receptive, individual test flowers were pollinated by depositing abundant pollen directly from the anthers of a pollen donor plant onto its stigma. Measures were taken for control and post-pollination test flowers in a staggered manner every two hours such that flowers of each plant were measured every four hours over a 12-hour day.

#### Model selection

We assessed the performance of machine learning model types commonly applied to spectral datasets: partial least squares (PLS), support vector machine (SVM), K-nearest neighbours (KNN), and random forest (RF), combining each model type with each of four pre-processing methods: unprocessed spectra, probabilistic quotient normalisation, first and second derivatives. The trained models were challenged against the test datasets and their performance assessed by accuracy of prediction, sensitivity and specificity (for binary models predicting pollination status), or R-squared and root mean squared error (RMSE) (for regression models predicting time since pollination).

The best performing model was determined by specificity-sensitivity ratio (binary model) or RFE and R^2^ (regression model) and selected as the final model. The cost and gamma (regression) parameters were tuned by grid search with 25-fold cross-validation.

#### Permutation test

Due to the slightly unbalanced dataset, a permutation test was conducted to calibrate the performance of our models, given the interdependency of the repeated measures. The null hypothesis of the permutation test is that there is no difference between treatment groups (pollination status, or time since pollination). The permutation test was executed with a SVM on data prepared as for the final model, and using the same parameters.

#### Metabolomics methodology

Literature sources used for library construction

- Jia et al., 2021, Comparative transcriptomic and metabolomic analyses of carotenoid biosynthesis reveal the basis of white petal color in *Brassica napus*.
- Chen et al., 2018, De novo transcriptome assembly, gene expressions and metabolites for flower color variation of two garden species in Brassicaceae
- Fu et al., 2018, Production of red-flowered oilseed rape via the ectopic expressions of *Orychophragmus violaceus* OvPAP2
- Hao et al., 2022, BnaA03.ANS Identified by Metabolomics and RNA-seq Partly Played Irreplaceable Role in Pigmentation of Red Rapeseed (*Brassica napus*) Petal
- Liu et al., 2021, Cross-Species Metabolic Profiling of Floral Specialized Metabolism Facilitates Understanding of Evolutional Aspects of Metabolism Among Brassicaceae Species

### Supplementary Results

#### Initial data exploration

The number of flowers differed across each timeseries group (), due to two principal factors; i) flowers were removed from testing if they became damaged during handling, or when the petals naturally dropped during senescence, ii) flowers were added to the experimental protocol successively as they opened, meaning the testing was conducted in a staggered manner. A proportion of each flowers’ time series was therefore ‘overnight’, as testing was conducted between the hours of 8 am and 8 pm.

#### Model selection

We conducted an exploratory analysis to determine the model to be applied to our dataset (Supp. Figure *8*,). In both the binary classification and regression we found SVM to have the highest performance out of those we tested, although its performance was comparable to PLS approach. Conversely, it is logical that the performance of KNN was poor, as to obtain reliable local estimates with such high dimensionality, a very high sample size would be required, as has been observed elsewhere (Melgani & Bruzzone, 2004). KNN is therefore less commonly used than other model types in applied contexts (Sarić et al., 2022).

For binary classification the linear SVM performed better than any kernel SVM. This suggests the pollinated and unpollinated flower spectra are linearly separable, which is particularly likely for a binary classification. In the case of the regression model, RBF kernel SVM outperformed the linear model. The nature of the regression task suggests mapping the features to higher dimensionality may allow non-linear relationships to be captured. The flexibility of the RBF kernel thus is suited to the complexity of the regression task (Pal & Foody, 2010).

#### Permutation test

The outcome of the permutation test for both the binary classifier and regression task reflected our slightly unbalanced dataset, with an accuracy of 57% for the binary SVM The R^2^ of the regression SVM on permuted data was 0.02 (**Error! Reference source not found.**). A Wilcoxon Mann-Whitney test showed there was a significant difference between the distributions of permuted training performance and the cross-validated training performance (p = 2.498 × 10 ^−7^), (**Error! Reference source not found.**)

### Supplementary Figures

**Supp. Figure 1.**
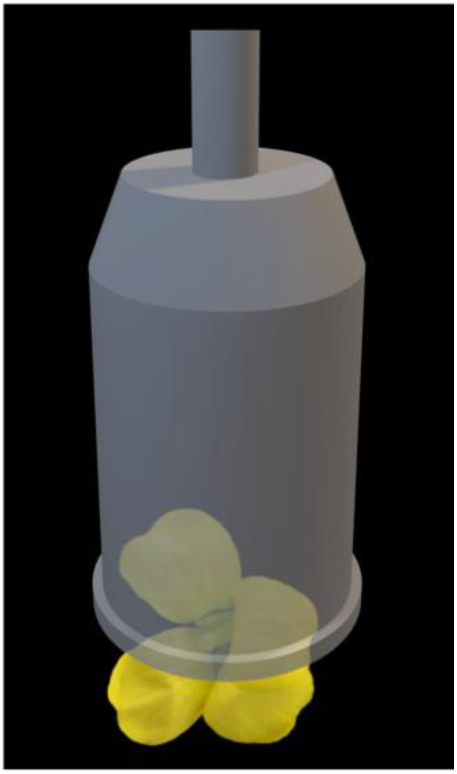
3D-printed probe mount fits over the flower being tested to ensure external light is excluded and only the full-spectrum light source illuminates the petal tissues.

**Supp. Figure 2.**
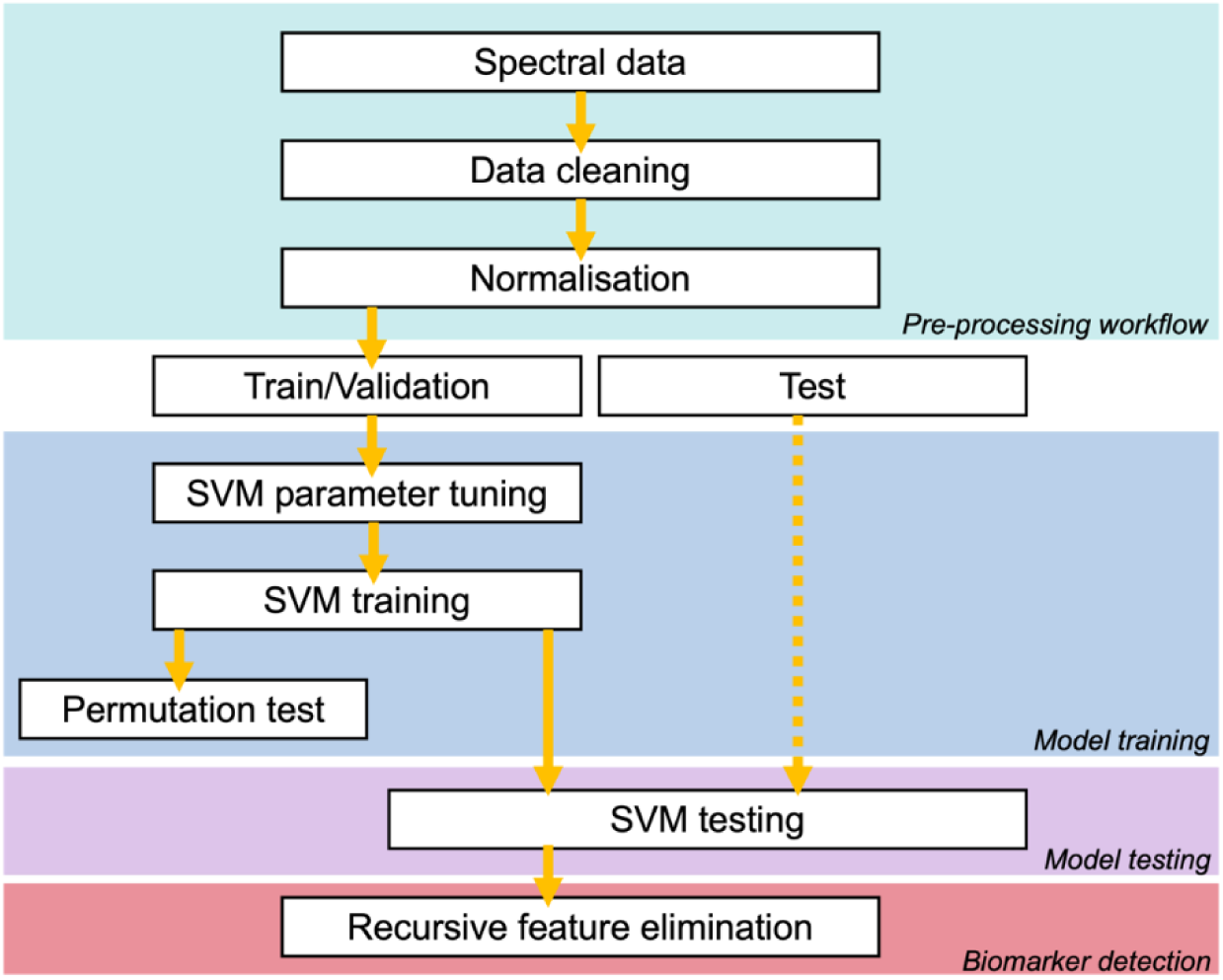
Summary of the final model pre-processing and model training and testing workflow.

**Supp. Figure 3.**
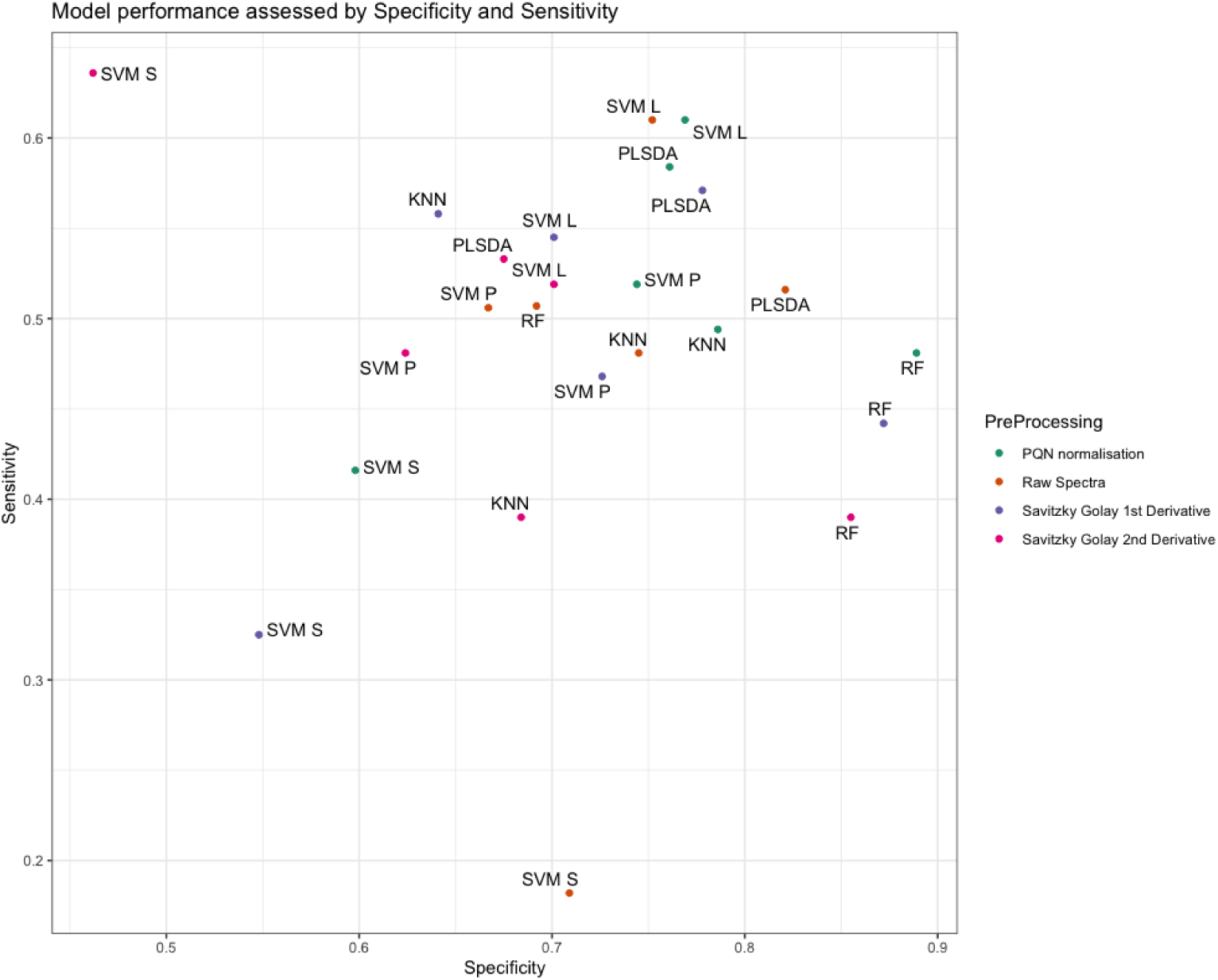
Model selection - Classification of pollination status by binary classification machine learning approaches, combined with pre-processing methods (raw spectra - no pre-processing) Probabilistic Quotient Normalisation (PQN), 1^st^ and 2^nd^ Derivatives calculated by Savitzky-Golay algorithm), assessed by classification sensitivity and specificity, with highest combined sensitivity and specificity being the best performing model.

**Supp. Figure 4.**
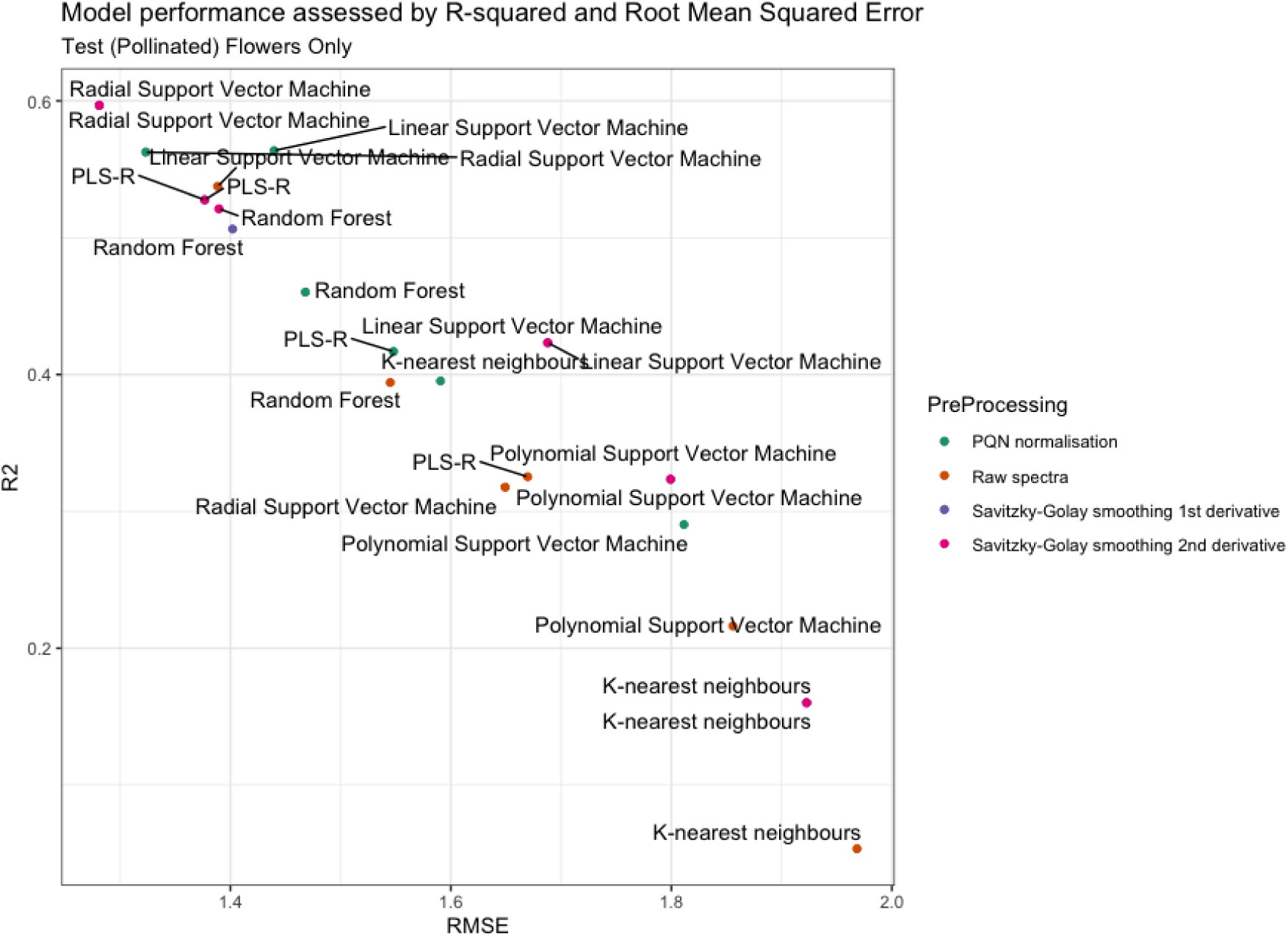
Model selection - Classification of post-pollination senescence by machine learning regression methods, combined with pre-processing methods (raw spectra - no pre-processing), Probabilistic Quotient Normalisation (PQN), 1^st^ and 2^nd^ Derivative (calculated by Savitky-Golay algorithm), assessed by R^2^ and Root Mean Squared Error (RMSE), with highest R^2^ and lowest RMSE being the best performing model.

**Supp. Figure 5.**
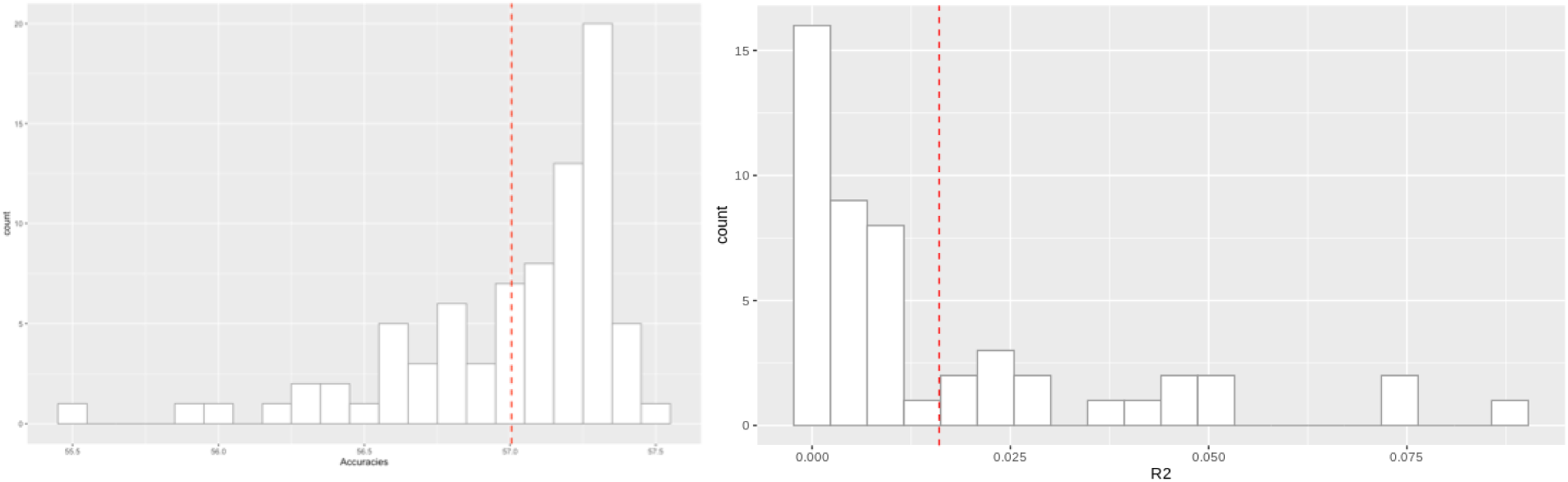
Result of the permutation test showing the performance of (left) the binary SVM predicting pollination status, assessed by accuracy, and (right) the regression SVM predicting time since pollination assessed by R^2^, on randomly permuted data.

**Supp. Figure 6.**
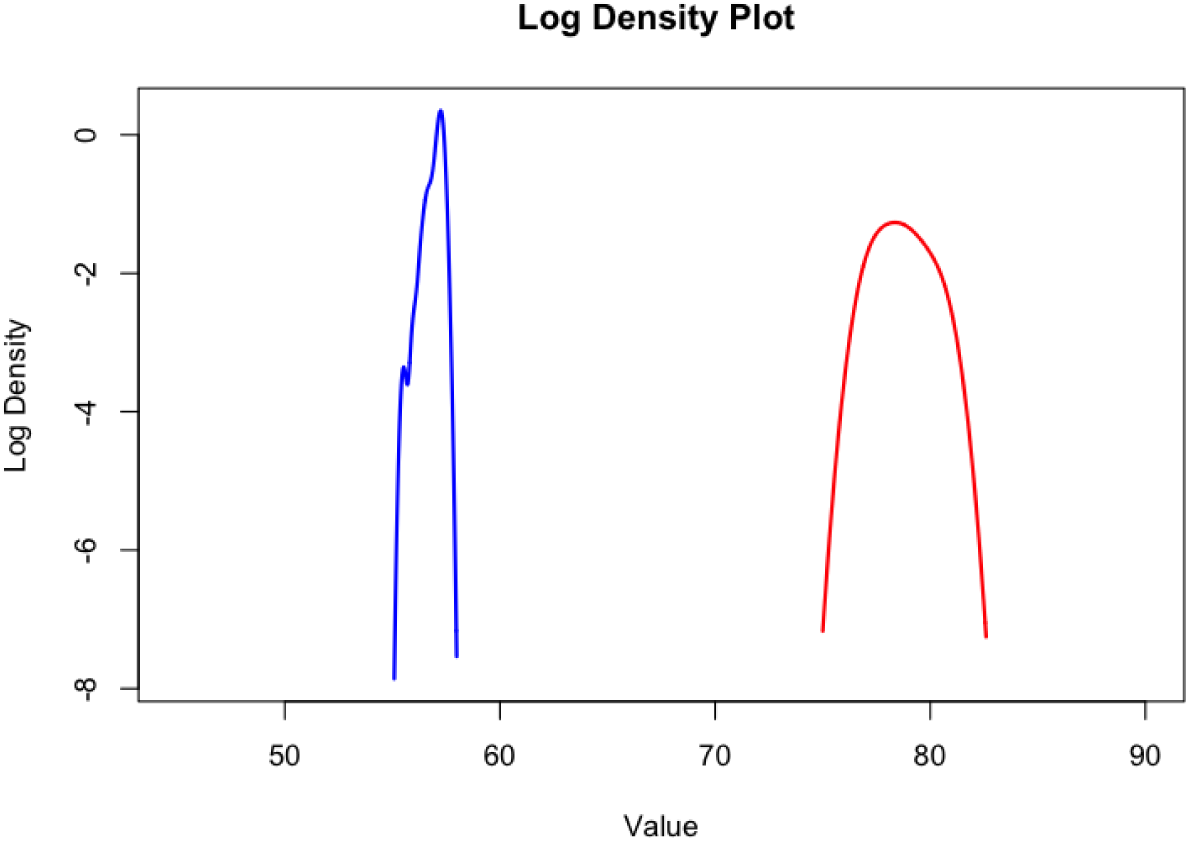
Distribution (natural log) of the permutation test accuracy (blue)

**Supp. Figure 7.**
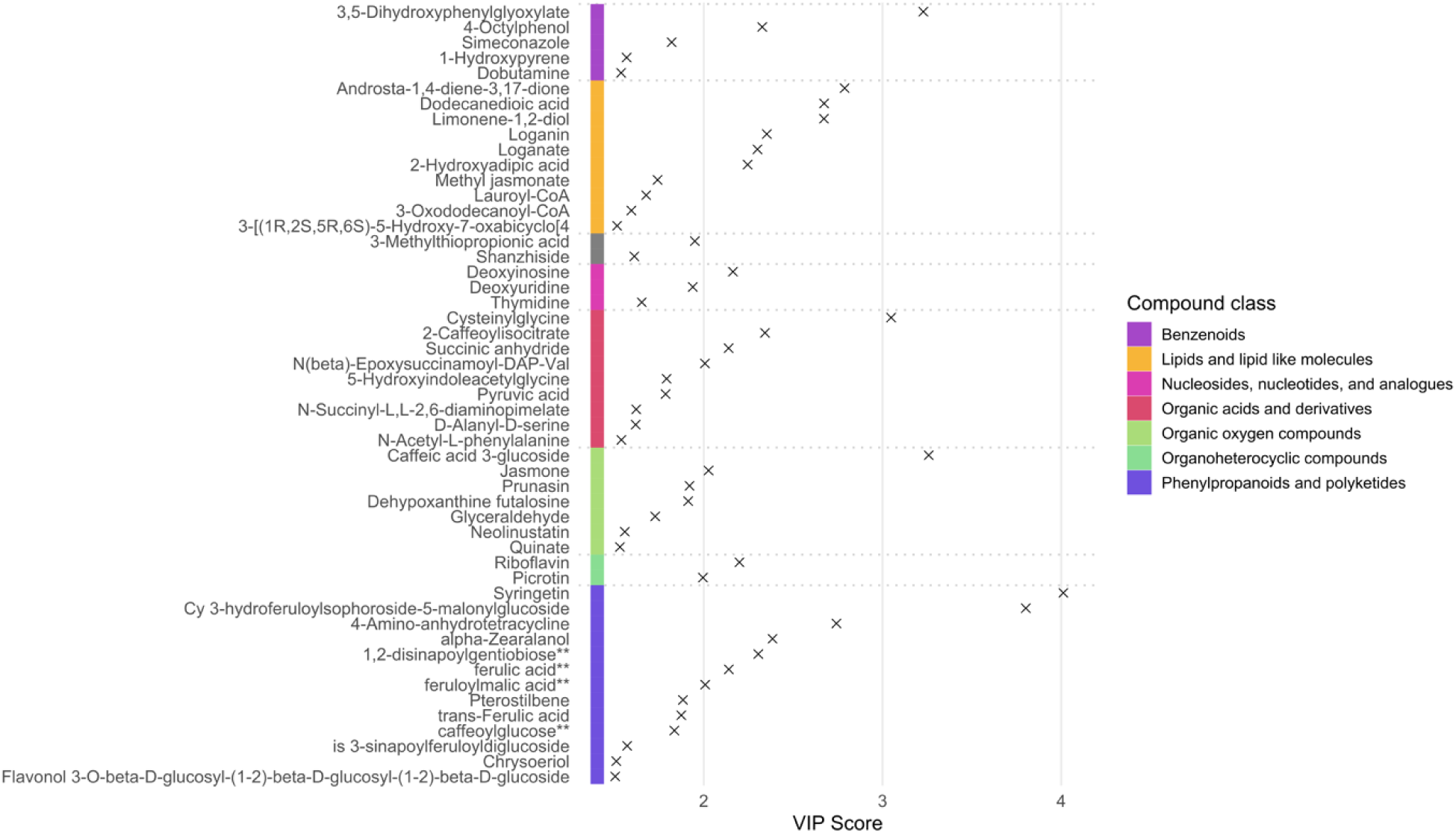
Compounds differentially expressed in pollinated and unpollinated flowers. Variable importance in projection (VIP) scores from the most important metabolites differential between pollination treatments (pollinated vs. unpollinated).

**Supp. Figure 8.**
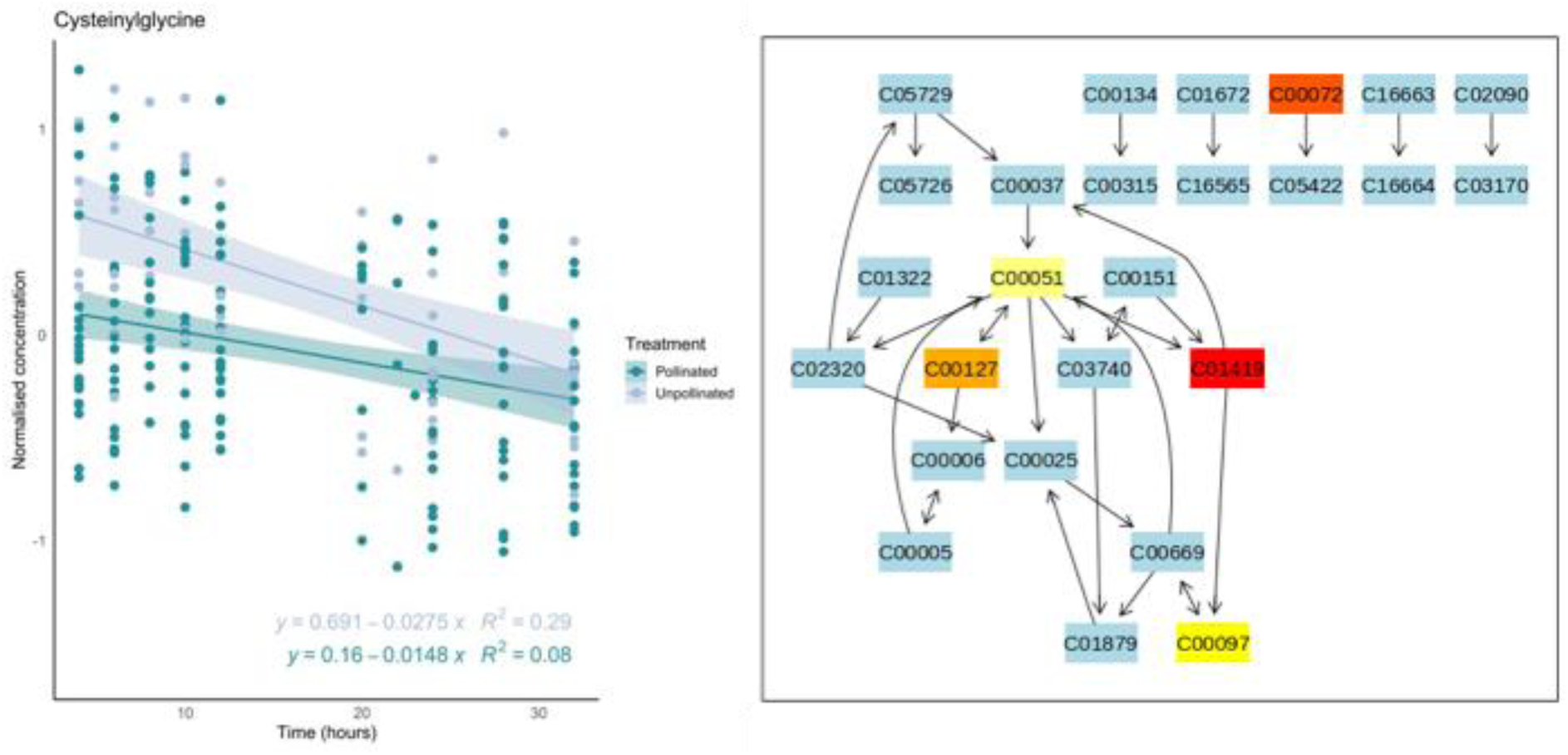
Cysteinylglycine and the glutathione metabolism pathway as an indicator of pollination. Left, normalised concentration of Cysteinylglycine (KEGGID C01419) over time between pollination treatment (pollinated vs. unpollinated. Right, KEGG Pathway for glutathione metabolism; blue indicates compounds that were not found in our metabolite samples, yellow-red scale indicates the significance of the metabolite to the pathway. C00051 – Glutathione, C00072 – Ascorbate, C00097 – L-Cysteine, C00127 – Glutathione disulfide, C01419 – Cysteinylglycine.

**Supp. Figure 9.**
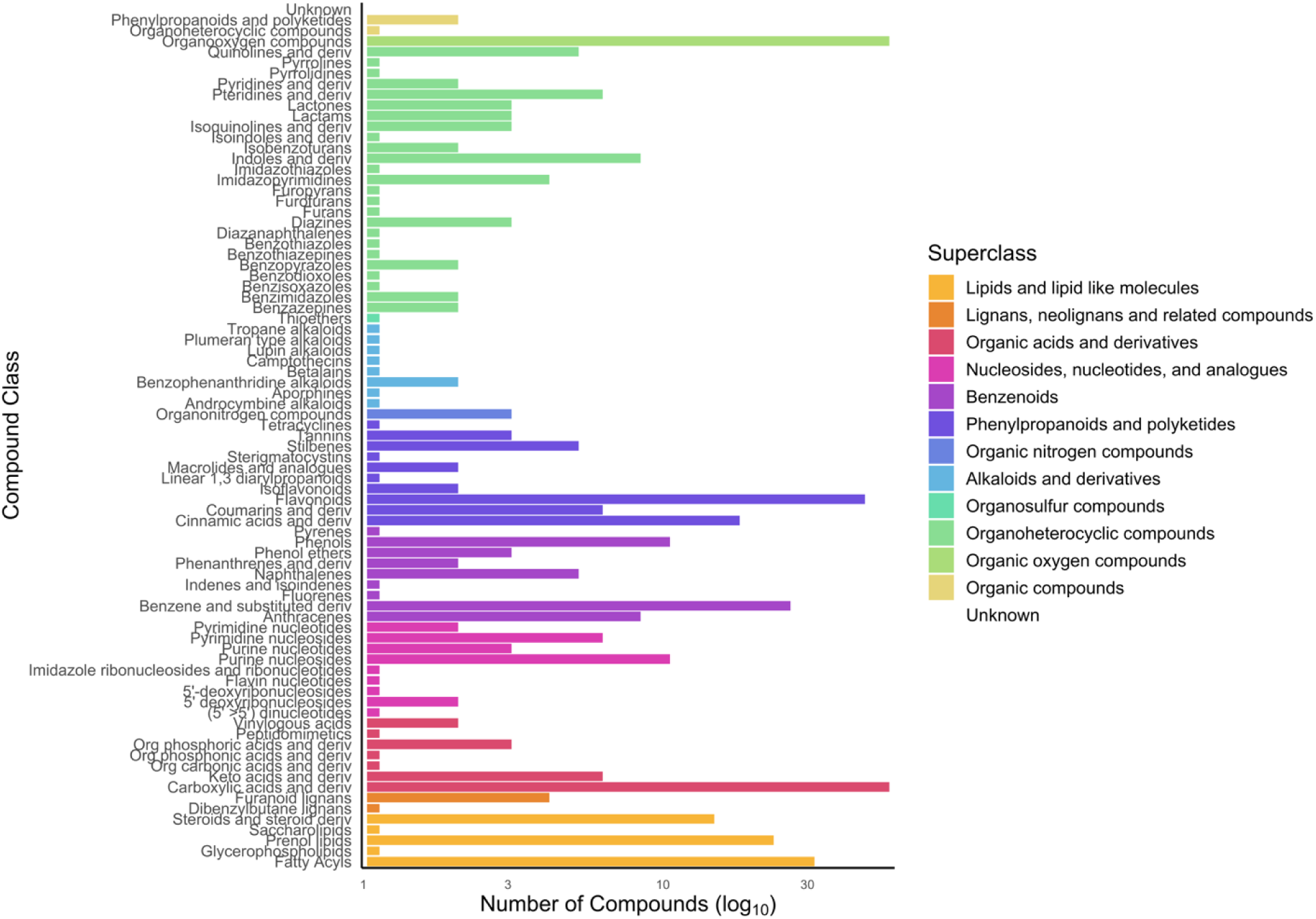
Frequencies of compound classes found in pollinated and unpollinated flowers over their development. Compound class frequencies (log_10_) grouped by their superclass derived from ChemOnt taxonomy.

**Supp. Figure 10.**
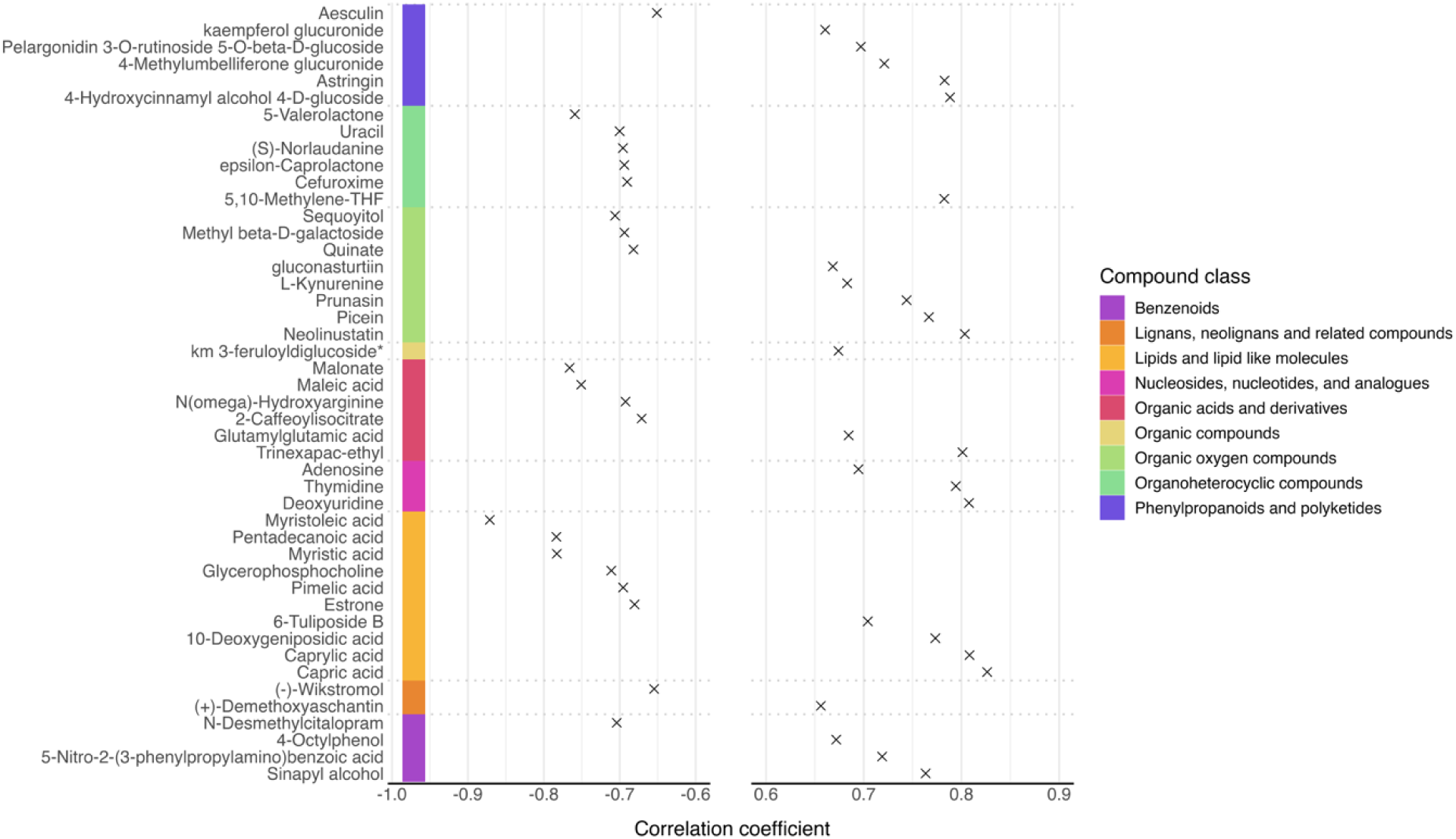
Compounds up- or down- regulated over time in pollinated and unpollinated flowers. Compounds most correlated (positive and negative Pearson’s correlation coefficients) with time, labelled by their taxonomical superclass (ChemOnt, (Djoumbou Feunang et al., 2016)).

### Supplementary Tables

**Supp. Table 1.** Sample size (number of flowers) grouped by time series (4 hour window from T1-8) and pollination status. T_0_ refers to pre-pollination measures.

|  | Time series | N = |
| --- | --- | --- |
| Unpollinated | T<sub>1</sub> | 63 |
|  | T<sub>2</sub> | 50 |
|  | T<sub>3</sub> | 45 |
|  | T<sub>4</sub> | 41 |
|  | T<sub>5</sub> | 38 |
|  | T<sub>6</sub> | 48 |
|  | T<sub>7</sub> | 29 |
|  | T<sub>8</sub> | 16 |
| Pollinated | T<sub>0</sub> | 75 |
|  | T<sub>1</sub> | 75 |
|  | T<sub>2</sub> | 69 |
|  | T<sub>3</sub> | 59 |
|  | T<sub>4</sub> | 53 |
|  | T<sub>5</sub> | 50 |
|  | T<sub>6</sub> | 48 |
|  | T<sub>7</sub> | 41 |
|  | T<sub>8</sub> | 22 |

